# Sub-Picomolar Detection of SARS-CoV-2 RBD via Computationally-Optimized Peptide Beacons

**DOI:** 10.1101/2021.06.04.447114

**Authors:** Soumya P. Tripathy, Manvitha Ponnapati, Joseph Jacobson, Pranam Chatterjee

## Abstract

The novel coronavirus SARS-CoV-2 continues to pose a significant global health threat. Along with vaccines and targeted therapeutics, there is a critical need for rapid diagnostic solutions. In this work, we employ deep learning-based protein design to engineer molecular beacons that function as conformational switches for high sensitivity detection of the SARS-CoV-2 spike protein receptor binding domain (S-RBD). The beacons contain two peptides, together forming a heterodimer, and a binding ligand between them to detect the presence of S-RBD. In the absence of S-RBD (OFF), the peptide beacons adopt a closed conformation that opens when bound to the S-RBD and produces a fluorescence signal (ON), utilizing a fluorophore-quencher pair at the two ends of the heterodimer stems. Two candidate beacons, C17LC21 and C21LC21, can detect the S-RBD with limits of detection (LoD) in the sub-picomolar range. We envision that these beacons can be easily integrated with on-chip optical sensors to construct a point-of-care diagnostic platform for SARS-CoV-2.

## Introduction

As numerous countries are experiencing additional waves of COVID-19, rapid, point-of-care diagnostic tests enable triage of symptomatic individuals and control the outbreaks of the disease. The most widely employed diagnostic tests for SARS-CoV-2 are reverse transcription-polymerase chain reaction (RT-PCR)-based methods (*1*), though other technologies based on CRISPR and loop-mediated amplification have been deployed as well (*2–5*). The best-in-class FDA authorized diagnostics, such as RT-PCR, have limits of detection (LoD) of 10^2^-10^3^ RNA copies/ml, which is about 1-10 attomolar (aM) RNA in the test volume (*6*). RT-PCR tests, however, require laborious and expensive nucleic acid isolation, purification, and processing steps, which increases both the turnaround time of detection and the cost of testing (*6, 7*). Alternatively, there are FDA-authorized low-sensitivity, inexpensive, and rapid diagnostics. These tests, which often rely on antigen detection, have LoDs of 10^5^-10^7^ RNA copies/ml, or around 1-100 femtomolar (fM) (*8*).

Recently, there has been significant effort to detect SARS-CoV-2 via fluorescence-based readouts to allow for specific signal amplification (*9–12*). Such methods largely rely on binding to SARS-CoV-2 RNA or DNA, which requires isolation of nucleic acids, as described above. In this study, we develop a molecular assay to detect the spike protein receptor binding domain (S-RBD) of SARS-CoV-2 using computationally-validated peptide beacons, which enable single-step detection of S-RBD presence through the production of a fluorescence signal. Our eventual goal is to integrate these optimized beacons within miniaturized total internal reflection fluorescence (TIRF) microscopes, which provide exquisite sensitivity by exciting fluorophores present within nanometer proximity of the device surface (*13*), producing high signal-to-background ratios and enabling rapid and ultra-sensitive detection of SARS-CoV-2.

## Results

### Engineering of Peptide Beacon Architecture

Our molecular beacon design includes two heterodimer-forming peptides, a binding ligand to the S-RBD, as well as a fluorophore-quencher pair at the terminal ends of the beacon (Figure 1A). This fluorophore-quencher pair induces fluorescence quenching through the mechanism of Förster resonance energy transfer (FRET), where the efficiency of energy transfer between the fluorophore and quencher is proportional to their spatial distance. Hence, a small change in spatial distance between the two beacon arms can drastically change the FRET efficiency, thus affecting the fluorescent quantum yield of the fluorophore. Here, we utilized a commonly-used fluorophore-quencher pair: fluorescein isothiocyanate (FITC) and [4-(N,N-dimethylamino)phenylazo] benzoyl (DABCYL), respectively. Upon binding to the target protein, the fluorophore and quencher are separated enough to observe an increase in fluorescence signal proportional to the amount of S-RBD present (Figure 1B).

**Figure 1:**
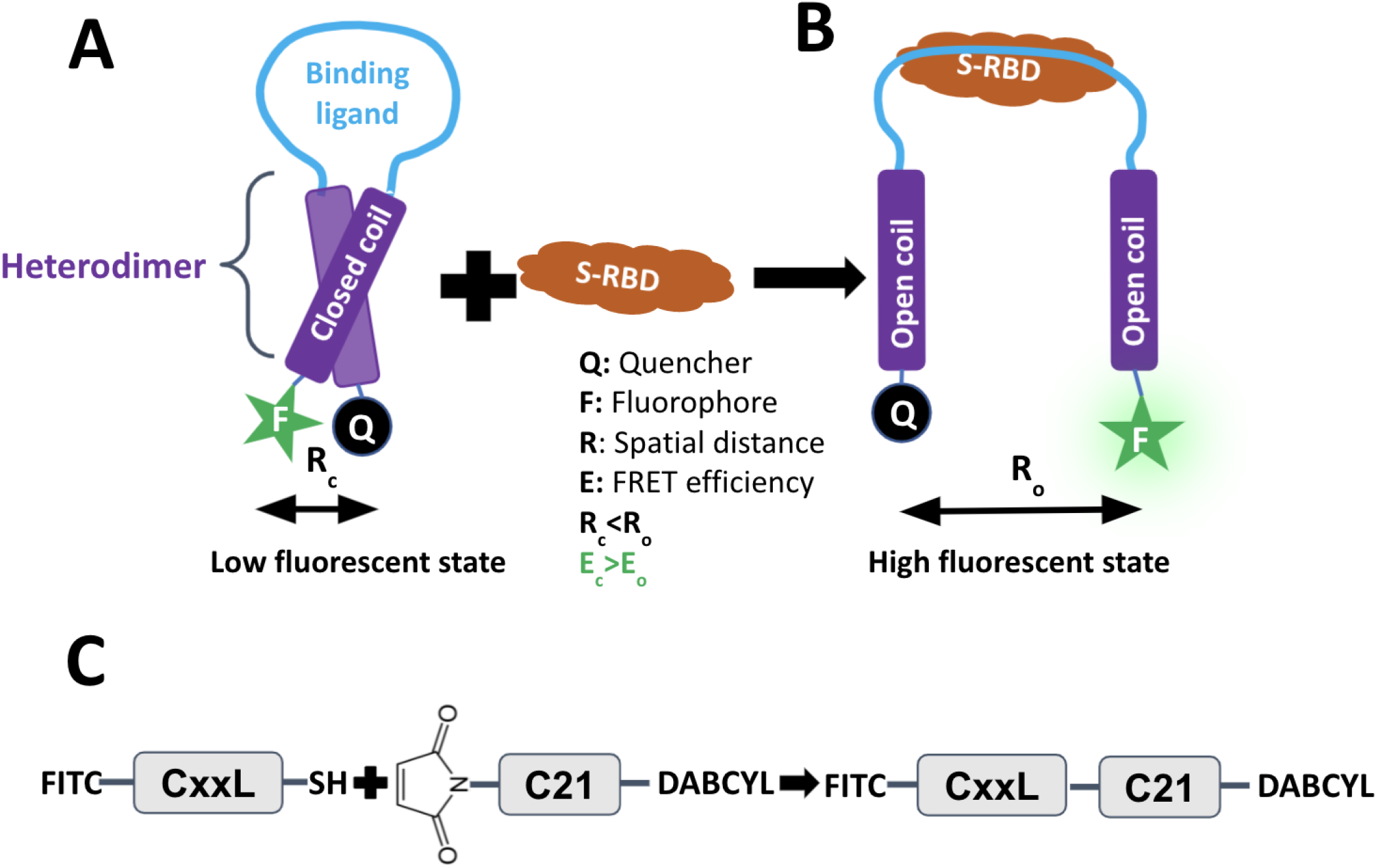
Engineering of peptide beacon architecture. A) Low-fluorescent state is the closed heterodimer state of the peptide beacon in the absence of S-RBD. B) High-fluorescent state is the open-coil state after binding of S-RBD with the loop of the peptide beacon. C) CxxL (25 uM, 1X PBS, pH 7.4) was added to C21 (25 uM, 1X PBS, pH 7.4) and incubated at room temperature for 2 hours for the conjugation reaction between the thiol groups of cysteine at the N-terminus of CxxL and the maleimide on the N-terminus of C21.

### Computational Selection of Peptide Beacon Candidates

We utilized the coiled-coil peptide beacons designed by Mueller, et al. (*14*). Mueller, et al., employed the parallel heterodimer reported by Thomas, et al., to design acidic 21mer peptide portions of a beacon detecting CREB binding protein (*15*). The reported peptide beacon designs consisted of the 21mer conjugated with a binding ligand and one of the three basic peptides: C13, C17, C21 (*14*). We designed our peptide beacons by replacing the binding portion of the sequences designed by Grossman with our previously engineered 23mer peptide that can bind to S-RBD and induce its degradation via the ubiquitin-proteasomal pathway (*16*).

To computationally verify that insertion of our 23mer peptide confers S-RBD binding capability to our peptide beacon sequences, we folded and docked the three designs (C13LC21, C17LC21, C21L21) using trRosetta, Rosetta, and HDOCK (Figure 2A) (*17–19*). Our results show that in the absence of S-RBD, all three peptides show terminal ends of the beacons, representing the fluorophore and the quencher, in close proximity to each other (OFF) in at least one of the top predicted models from tr-Rosetta (*17*) or Rosetta Abinitio (*18*) (Figure 2B-2C). Alternatively, when the peptide beacon sequences were docked against S-RBD utilizing HDOCK (*19*), we observed the terminal ends to be spatially distanced from each other, indicating a possible ON state (Figure 2D). These results motivated us to test these three peptide beacon designs experimentally.

**Figure 2:**
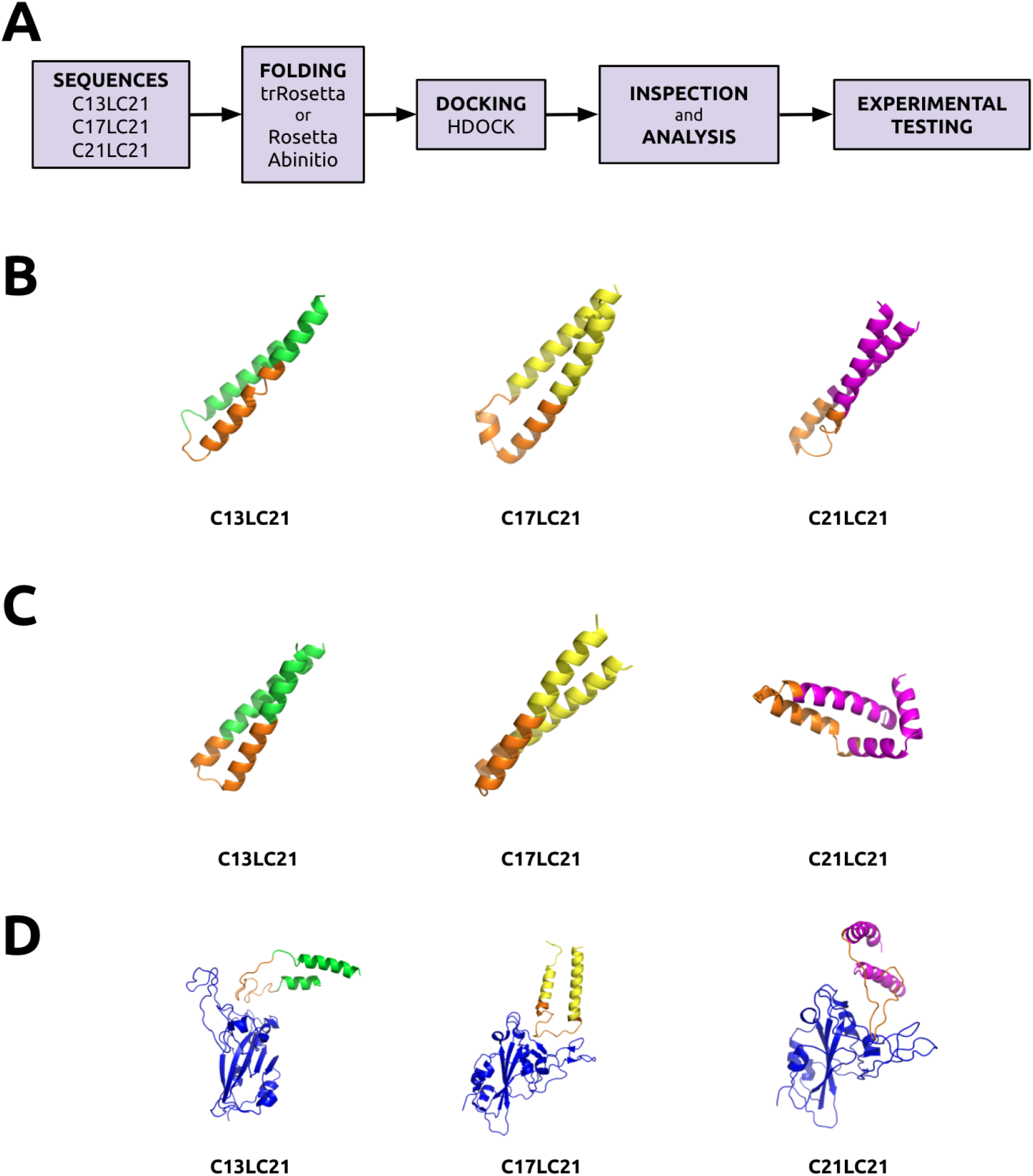
*In silico* verification of peptide beacon architecture. A) Validation pipeline. Sequences were folded ab initio via trRosetta or Rosetta Abinitio. The top structures were docked to S-RBD via HDOCK. Docked structures were analyzed and candidate beacons were chosen. Highest scoring B) trRosetta structures C) Rosetta Abinitio structures and D) Docked HDOCK structures of the three candidate beacons. The conserved binding peptide moiety is highlighted in orange. S-RBD (derived from PDB 6MOJ) is indicated in blue.

### Validation of S-RBD Binding in Human Cells

To rapidly validate the binding capability of our peptide beacon designs, we adapted our previously-described degradation assay in human cells, by fusing our peptide beacon candidates to the CHIPΔTPR E3 ubiquitin ligase, which can tag target proteins for degradation via the ubiquitin-proteasomal pathway in human cells. (*16*). After co-transfection with a plasmid expressing S-RBD C-terminally fused to superfolder GFP (sfGFP), we can measure binding affinity to S-RBD as relative to sfGFP degradation. We tested various peptide beacon combinations, employing our validated mutant S-RBD-binding peptide derived from ACE2, A2N (*16*), as well as a positive nanobody control, which has shown high affinity to S-RBD (*20*). As expected, these moieties alone demonstrate robust degradation capabilities, while the arms-only negative controls (C13, C17, C21, and C21*) show negligible degradation, hence no binding to the S-RBD. Of the complete peptide beacon constructs, the C17-A2N-C21* (C17LC21) beacon induces the greatest level of degradation of S-RBD-sfGFP, followed by C21LC21 and C13LC21, the latter of which exhibited minimal degradation capabilities (Figure 3A).

**Figure 3:**
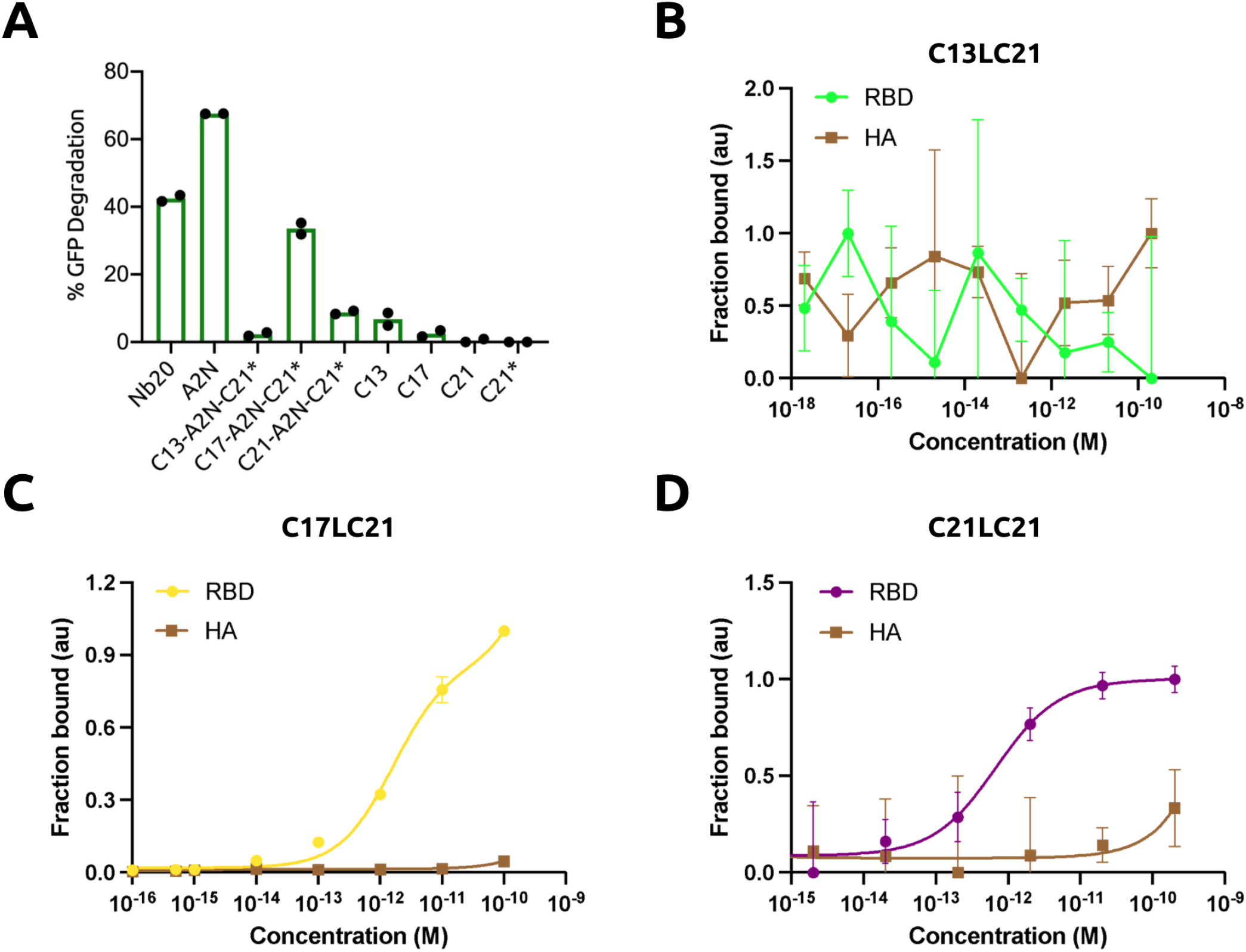
Experimental characterization of peptide beacons. A) Analysis of S-RBD-sfGFP degradation by flow cytometry. All samples were performed in independent transfection duplicates (n=2) and gated on GFP+ fluorescence. Mean percentage of GFP+ cell depletion was calculated in comparison to the S-RBD-sfGFP only control. C13LC21 is referred to as C13-A2N-C21*, C17LC21 is referred to as C17-A2N-C21*, and C21LC21 is referred to as C21-A2N-C21*. Titration of the target recombinant S-RBD and Influenza H3N2 (HA) with 2 nM of HPLC-purified B) C13LC21 (n=3), C) C17LC21 (n=5), D) C21LC21 (n=3) in 1X PBS, pH 7.4.

### *In vitro* Detection of S-RBD

After ascertaining the binding capability of our candidates in human cells, we characterized the response of the peptide beacons *in vitro* in the presence of S-RBD. As a negative control, we also measured the cross reactivity of our peptide beacons towards Hemagglutinin protein of Influenza A H3N2 (HA) (*21*). We first purified peptide beacons (CxxLC21) from the reaction mixture of CxxL+C21, using high performance liquid chromatography (HPLC) and confirmed the presence of CxxLC21 in the collected fraction from HPLC through MALDI-TOF mass spectrometry (Supplementary Figure 1). We measured the fluorescent signals from the peptide beacons following a 10 minute exposure to different concentrations of S-RBD and HA in 1X PBS and pH 7.4 (Figure 3B-D). Of the three peptide beacons, C17LC21 showed highest sensitivity towards S-RBD with a LoD of nearly 20 fM (*K*_*d*_ = 1.615 *×* 10^*-*12^), followed by C21LC21 having an LoD of 400 fM (*K*_*d*_ = 6.766 *×* 10^*-*13^). However, C13LC21 showed negligible response towards both S-RBD and HA. In conclusion, C17LC21 and C21LC21 are able to detect the presence of S-RBD with sub-picomolar sensitivity and low cross-reactivity, thus motivating their application for rapid detection of SARS-CoV-2.

## Discussion

In this study, using existing deep learning tools for protein structure prediction and energy-based modeling suites, we designed and tested a set of molecular beacons that can potently bind to the S-RBD and release a fluorescence signal via FRET, enabling sub-picomolar detection levels. Integration of these peptide beacons within optical sensors, such as miniature TIRF microscopes, may reduce the LoD to sub-femtomolar level, thus yielding a rapid, point-of-care diagnostic platform for SARS-CoV-2. Such a diagnostic would be faster than existing RT-PCR assays and more sensitive than antigen-based rapid testing.

Our pipeline also showcases a use case for current deep learning tools for protein structure prediction in a protein design pipeline. Using this approach, molecular beacons can be designed to detect protein targets within other viral species. In addition, by using existing protein structure prediction tools to get rapid insights into the structure of the protein, it may be possible to design an entire peptide beacon sequence from scratch even in the absence of a known binding partner. By fixing the heterodimer motif, binding loops of molecular beacons can be filled in via protein hallucination, using tools like tr-Rosetta (*22*). Thus, by employing a hybrid approach of state-of-the-art protein modeling tools and robust experimental validation, our molecular beacon design pipeline serves as a powerful platform to fight COVID-19 and future emergent viral threats.

## Materials and Methods

### *In silico* Selection of Candidate Peptide Beacons

The three peptide beacon sequences were folded in the absence of S-RBD using trRosetta. trRosetta is a deep learning tool to predict structures from sequence information (*17*). The three peptide beacon sequences were also folded using Rosetta Abinitio folding. Abinitio folding solves the protein structure from sequence through physics-based constraints rather than relying on previously solved structures like trRosetta (*18*).

The three peptide beacon sequences were docked against the S-RBD using HDOCK. HDOCK is a protein-protein docking platform that combines ab-initio docking and template-based modeling (*19*). The top 100 predictions from HDOCK were analyzed to visualize the structure of peptide beacon sequences in the presence of S-RBD.

### Generation of Plasmids

pcDNA3-SARS-CoV-2-S-RBD-sfGFP (Addgene #141184) and pcDNA3-R4-uAb (Addgene #101800) were obtained as gifts from Erick Procko and Matthew DeLisa, respectively. Peptide beacon sequences were ordered as gBlocks (IDT), and were amplified with overhangs for Gibson Assembly-mediated insertion into linearized pcDNA3-R4-uAb digested with HindIII and EcoRI. Assembled constructs were transformed into 50 *µ*L NEB Turbo Competent *E. coli* cells, and plated onto LB agar supplemented with the appropriate antibiotic for subsequent sequence verification of colonies and plasmid purification.

### Cell Culture

HEK293T cells were maintained in Dulbecco’s Modified Eagle’s Medium (DMEM) supplemented with 100 units/mL penicillin, 100 mg/mL streptomycin, and 10% fetal bovine serum (FBS). RBD-sfGFP (250 ng) and peptide-E3 ligase fusion (250 ng) plasmids were transfected into cells (2 × 10^5^/well in a 24-well plate) with Lipofectamine 3000 (Invitrogen) in Opti-MEM (Gibco). After 5 days post-transfection, cells were harvested and analyzed on a BD FACSCelesta flow cytometer (BD Biosciences) for GFP fluorescence (488-nm laser excitation, 530/30 filter for detection). Cells expressing GFP were gated, and percent GFP+ depletion to the RBD-sfGFP only control were calculated. All samples were performed in independent transfection duplicates (n=2), and percentage depletion values were averaged. Standard deviation was used to calculate error bars.

### Peptide Synthesis and Purification

CxxL (25 *µ*M, 1X PBS, pH 7.4) was added to C21 (25 *µ*M, 1X PBS, pH 7.4) and incubated at room temperature for 2 hours for the conjugation reaction between the thiol group of Cysteine at N-terminus of CxxL and the maleimide group at the N-terminus of C21 (Figure 1C). CxxL+C21 is used to represent the reaction mixture obtained after 2 hours of reaction between CxxL and C21 in 1X PBS, pH 7.4 at room temperature. MALDI-TOF mass spectroscopy of CxxL+C21 confirms the presence of CxxLC21 in CxxL+C21 along with CxxL and C21 (Supplementary Figure 1A-1C). The reaction mixture of CxxL and C21 contains the mixture of individual CxxL, C21, and CxxLC21 after 2 hours of reaction in 1X PBS, pH 7.4 at room temperature. Consequently, CxxLC21 from CxxL+C21 was purified using high performance liquid chromatography (HPLC) (Agilent 1100) using a C14 H31 column. HPLC chromatogram of CxxL+C21 contains peaks at distinct retention time than that of CxxL and C21 in HPLC chromatogram (Supplementary Figure 1D-1F), which are considered as the fraction containing CxxLC21. The fraction was collected and MALDI-TOF mass spectroscopy was performed (Microflex LRF, Bruker), confirming the presence of CxxLC21 at higher concentration than that of CxxL and C21 (Supplementary Figure 1G-1I). The collected fraction was freeze-dried and used for detection of S-RBD and HA.

### *In vitro* Detection of S-RBD

S-RBD (Abcam ab273065) and Influenza H3N2 hemaglutinin (HA) (MyBioSource MBS5308351) were titrated against purified CxxLC21. S-RBD and HA proteins were serially diluted in 1X PBS, pH 7.4 from nanomolar (nM) to attomolar (aM) concentration. 2 nM of CxxLC21 in 1X PBS, pH 7.4 (n=3 or 5) was exposed to different concentrations of the target protein (S-RBD or HA) and incubated for 10 minutes at room temperature. Fluorescence intensity was subsequently measured using a Tecan Spark well plate reader at excitation and emission wavelength of 470 nm and 525 nm, respectively. The fluorescence data was fitted to one site total binding with saturation curve in the Prism software using the following equation:

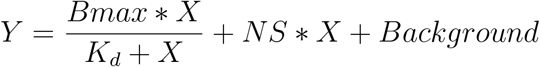

where *Y* is the fraction of peptide beacon bound to target, *Bmax* is the maximum binding in units of *Y, X* is the concentration of target, *K*_*d*_ is the equilibrium dissociation constant in units of *X*, and *NS* is the slope of the non-linear regression. The fraction of peptide beacon bound to target (*Y*) is calculated by normalizing fluorescent intensities obtained at different (*X*) by considering highest fluorescence intensity obtained at maximum S-RBD concentration as hundred percent. The limit of detection (LoD) is estimated from the equation given below.

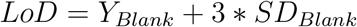

### Statistics and Reproducibility

All samples were performed in independent duplicates (n=2), triplicates (n=3), or quintuplicates (n=5), as indicated. Standard deviation was used to calculate error bars. Statistical analyses was performed using the two-tailed Student’s t test, using the GraphPad software package.

## Acknowledgments

We thank Dr. Neil Gershenfeld, Dr. Shuguang Zhang and Dr. Edward S. Boyden for shared lab equipment, and Teodora Stan for techical assistance. This work was supported by the consortia of sponsors of the MIT Media Lab and the MIT Center for Bits and Atoms, and by Jeremy and Joyce Wertheimer.

## Competing Interests

P.C., M.P., S.P.T and J.M.J. are listed as inventors for U.S. Provisional Patent Application 63/182,537 entitled “Peptide Based Probes For the Detection of SARS-CoV-2.”

## Author Contributions

S.P.T. conducted peptide beacon purification, synthesis, and *in vitro* characterization. M.P. conducted computational design and docking protocols, and performed experiments. P.C. built constructs, performed degradation assays, and conducted data analysis. P.C., S.P.T, and M.P. wrote the manuscript. P.C. and J.M.J supervised the project.

## Data and Code Availability Statement

All data needed to evaluate the conclusions in the paper are present in the paper. All source computational and experimental data files can be accessed at: https://tinyurl.com/covidpepbeacons.

---

**Figure S1:**
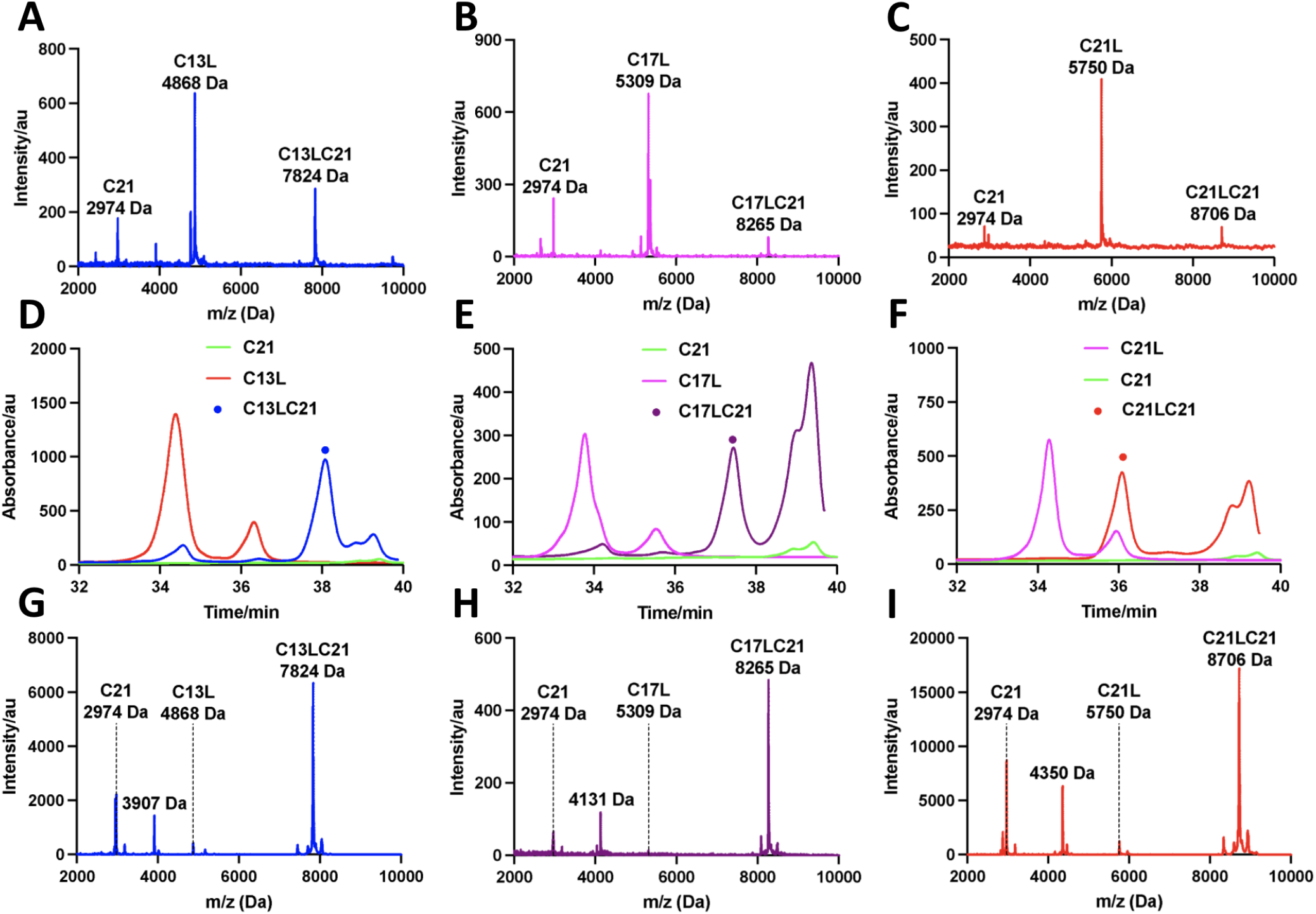
Purification and synthesis of peptide beacons. A-C) MALDI-TOF mass spectrum of CxxL+C21 shows peaks corresponding to C21, CxxL, and CxxLC21. D-F) HPLC chromatograms of CxxLC21, plotted with HPLC chromatograms of CxxL and C21. The chromatogram of C13L+C21, C17L+C21, and C21L+C21 shows the appearance of new peaks having retention times of 38 min, 37 min, and 36 min, respectively (new peaks marked in colored dot in D-E). Fraction of material corresponding to new peaks was collected during HPLC. G-I) MALDI-TOF mass spectrum of collected fraction from HPLC of CxxL+C21 shows dominant peak corresponding to CxxLC21 and negligible peaks corresponding to CxxL and C21. CxxL+C21 represents the reaction mixture of CxxL and C21 after 2 hours of reaction at room temperature. CxxLC21 represents the peptide beacons.

## Notes

https://tinyurl.com/covidpepbeacons

